# Hierarchical Matrix Compression for Fast Bidomain Simulation of Multicellular Neuron Models

**DOI:** 10.64898/2026.11.20.768225

**Authors:** Rodrigo Esparza Rivera, Luis J. Gomez

**Affiliations:** Elmore Family School of Electrical and Computer Engineering, Purdue University, West Lafayette, Indiana, USA

## Abstract

The bidomain boundary element–cable method (Cable-BEM) couples the intracellular, membrane, and extra-cellular dynamics of multicellular neuron models using only the compartmental unknowns of cable models. However, its matrices are dense, and their storage and factorization scale as *O*(*N* ^2^) and *O*(*N* ^3^) for *N* unknowns, respectively. We compare four hierarchical matrix formats for compressing these matrices: hierarchically off-diagonal low-rank (HODLR), hierarchically semiseparable (HSS), *H*-, and *H*^2^-matrices. For a layer 2/3 pyramidal neuron, the four formats use 2.7 to 6.1 times less memory than the dense solver. At a tolerance of 10^*−*3^, the maximum membrane potential errors are 8.5 mV for HSS, 0.47 mV for HODLR, and at most 10^*−*3^ mV for the *H*- and *H*^2^-matrices. For arrays of 9 to 225 parallel axons (1,827 to 45,675 unknowns), the maximum rank reaches 1,759 for HSS and 797 for HODLR, whereas it remains at or below 101 for the *H*- and *H*^2^-matrices. These results suggest that *H*- and *H*^2^-matrices scale more favorably than HODLR and HSS for networks of closely packed cells.

## I. Introduction

Extracellular electric fields influence neural activity through externally applied stimulation and endogenous ephaptic coupling. Cable models typically prescribe the extracellular potential and therefore do not capture the feedback between membrane currents and the extracellular field [1]. Fully coupled finite element models capture this feedback, but they require volumetric meshes that are computationally expensive for realistic multicell geometries [2]. Cable-BEM is a wire-kernel bidomain boundary element method that retains the compartmental unknowns of cable models while coupling the intra-cellular and extracellular fields [3]. It integrates the boundary integral kernels around each cylindrical compartment and thus avoids meshing the membrane surfaces. However, the resulting matrices are dense. For *N* unknowns, they require *O*(*N* ^2^) memory, and a dense factorization requires *O*(*N* ^3^) operations.

In [3], the memory was reduced by a factor of up to 4.6 for a 225-cell network using adaptive cross approximation (ACA) and hierarchically off-diagonal low-rank (HODLR) compression. HODLR uses weak admissibility; i.e., it compresses every off-diagonal block regardless of the distance between the corresponding groups of unknowns. Blocks that couple neighboring compartments or cells can require high ranks, which limits the compression as the network grows.

Here, we compare HODLR with hierarchically semiseparable (HSS) matrices and with strongly admissible *H*- and *H*^2^-matrices for compressing the Cable-BEM matrices. We report the accuracy, memory, run time, and maximum rank of each format for a single neuron and for arrays of up to 225 axons.

## II. Bidomain Boundary-Element-Cable Formulation

Cable-BEM represents each neuron as a graph of tapered cylindrical compartments. Each node carries one unknown for the transmembrane voltage *V*_*m*_ and one for the interface charge density *ρ*. Under the quasi-static approximation, enforcing continuity of the normal conduction current across all membrane interfaces yields the bidomain boundary integral equation, which is advanced in time using a semi-implicit Crank–Nicolson scheme. The azimuthal dependence of the cylindrical kernels is integrated analytically using elliptic integrals, which reduces all source integrals to one-dimensional axial quadrature. Each time step requires the solution of

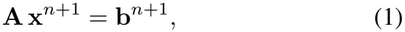

where **A, x**^*n*+1^, and **b**^*n*+1^ are the dense *N × N* system matrix, the unknowns at time step *n* + 1, and a right-hand side that contains the membrane state and the device forcing, respectively, and *N* is the number of compartments in all cells. Because **A** depends only on the geometry, it is compressed and factorized once and reused at every time step. Details of the formulation are given in [3].

## III. Hierarchical Matrices

Hierarchical matrices compress **A** by recursively grouping the unknowns by spatial proximity using a k-d tree. Each resulting matrix block is stored either densely or in low-rank form, as determined by the admissibility criterion of the format [4].

### A. Weak Admissibility

Under weak admissibility, every off-diagonal block is approximated in low-rank form regardless of the distance between the corresponding groups, and only the diagonal leaf blocks are stored densely. HODLR and HSS matrices use this criterion. In our HODLR implementation, ACA constructs the low-rank factors without assembling the off-diagonal blocks. HSS matrices additionally use nested bases; i.e., the basis of a parent cluster is constructed from the bases of its children through transfer matrices, which reduces the storage when the ranks are small.

### B Strong Admissibility

In a strongly admissible *H*-matrix, the block associated with clusters *τ* and *σ* is approximated in low-rank form only if dist(*τ, σ*) *≥ η* min {diam(*τ*), diam(*σ*)}, where *η >* 0 is the admissibility parameter. All other blocks are subdivided further or, at the leaf level, stored densely. An *H*^2^-matrix additionally uses nested cluster bases for its admissible blocks.

## IV. Results

We first simulate a rat layer 2/3 pyramidal neuron morphology from ModelDB [5]. A 200 pA current is injected into the axon, and *V*_*m*_ is recorded at the axon, soma, and apical dendrite along the path shown in Fig. 1(a). The simulation spans 14 ms in 2,000 time steps. The compressed matrices are constructed to a tolerance of 10^*−*3^, and the dense Cable-BEM solution is the reference. We use STRUMPACK [6] for the HSS and HODLR solvers, with ButterflyPACK [7] providing the HODLR format, and H2Lib [8] for the *H*- and *H*^2^-matrix solvers. All simulations were run on a single cluster node (2× AMD EPYC 9654, 96 cores each, 384 GB RAM), using 12 cores and 16 GB of memory per job. The code is MATLAB R2024a with C++ MEX kernels compiled with GCC 14.1 and parallelized with OpenMP.

**Fig. 1.**
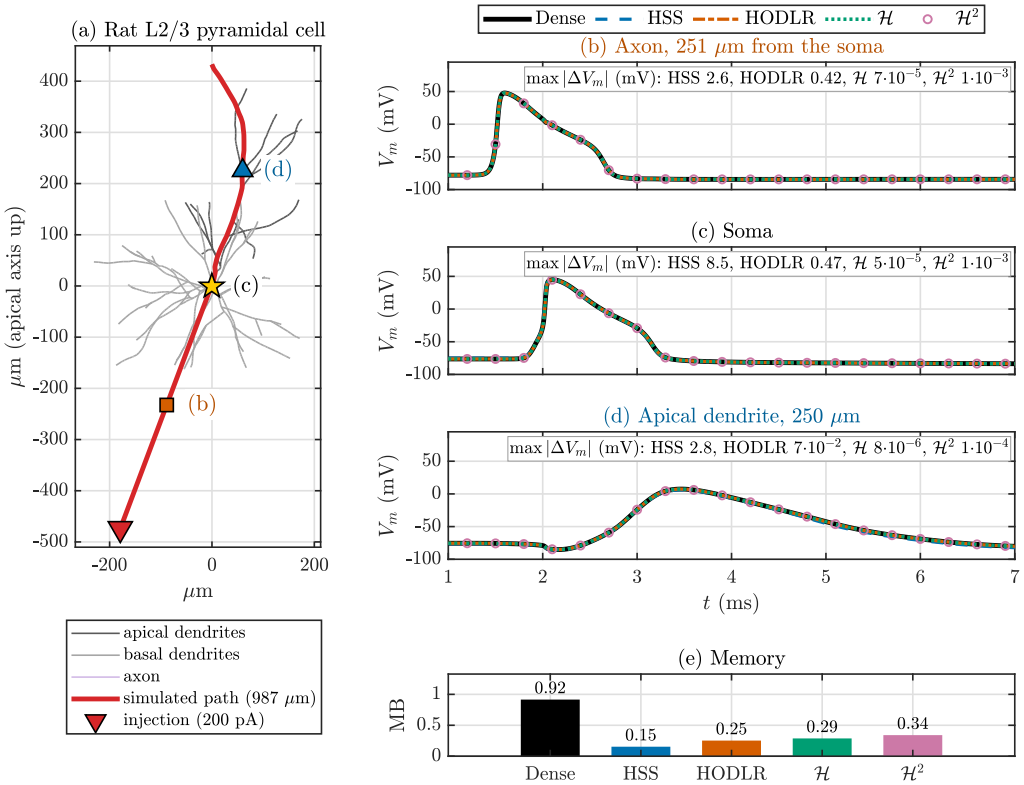
Single layer 2/3 pyramidal neuron. (a) Morphology, simulated path, and recording sites. (b)–(d) *V*_*m*_ at the axon, soma, and apical dendrite for the dense and compressed solvers (tolerance 10^*−*3^); each panel lists the maximum errors relative to the dense solution. (e) Memory for the matrices and their factors.

The membrane potentials are shown in Fig. 1(b)–(d). The maximum errors over the three sites are 8.5 mV for HSS, 0.47 mV for HODLR, 10^*−*3^ mV for ^2^, and 7 *×* 10^*−*5^ mV for *H*. Thus, the strongly admissible formats are over two orders of magnitude more accurate at the same nominal tolerance. The HSS error of 8.5 mV suggests that HSS requires a tighter tolerance than 10^*−*3^ for this problem. HSS, HODLR, *H*, and *H*^2^ use 6.1, 3.7, 3.2, and 2.7 times less memory for the matrices and their factors than the dense solver, respectively [Fig. 1(e)].

Next, we simulate square arrays of 3 *×*3, 5 *×*5, 7 *×*7, 10 *×* 10, and 15 *×* 15 parallel cylindrical axons with 203 unknowns each (1,827 to 45,675 unknowns) at a tolerance of 10^*−*5^ (Fig. 2). For 225 axons, each format requires less than 10 GB, whereas the dense solver, which was run for up to 100 axons, would require about 47 GB by extrapolation. The fastest (*H*) and slowest (HSS) compressed solvers take about 18 and 42 min for the 2,000 time steps, respectively, compared with a projected 84 min for the dense solver. The maximum rank grows as *N* ^1.05^ for HSS and *N* ^0.78^ for HODLR, reaching 1,759 and 797 for 225 axons, respectively. In contrast, it grows as *N* ^0.29^ for *H* and *N* ^0.28^ for *H*^2^, reaching 77 and 101, respectively. The rank growth of HSS and HODLR is likely due to the blocks that couple neighboring axons, which weak admissibility compresses regardless of their separation.

**Fig. 2.**
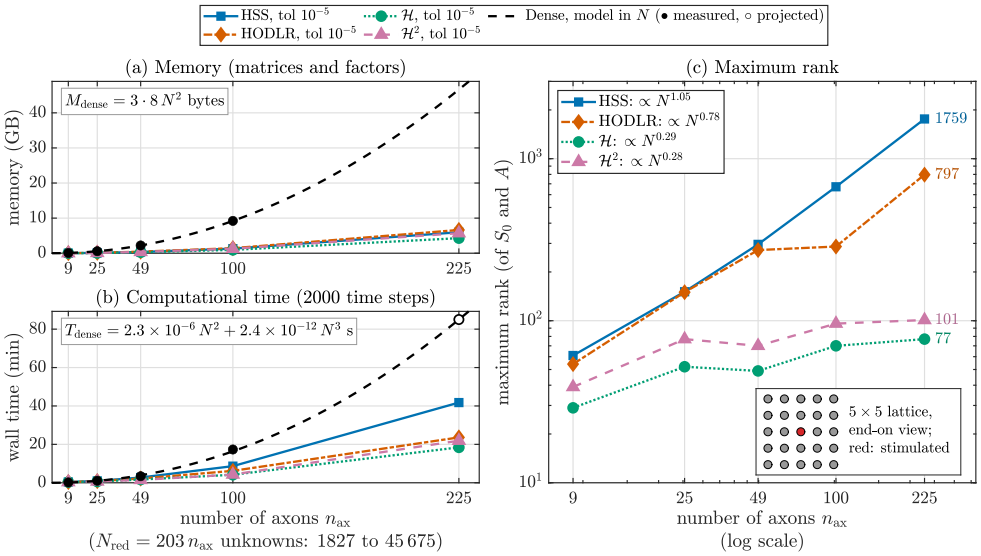
Arrays of 9 to 225 parallel cylindrical axons at a tolerance of 10^*−*5^. (a) Memory for the matrices and their factors. (b) Wall time for 2,000 time steps. (c) Maximum rank. The dense results are measured for up to 100 axons and extrapolated for 225 axons.

## V. Conclusion

We compared HODLR, HSS, *H*-, and *H*^2^-matrix compression of the Cable-BEM matrices. For a single neuron, the four formats reduce the memory by factors of 2.7 to 6.1, and the strongly admissible formats are the most accurate at a given tolerance. For arrays of up to 225 axons, the maximum rank of the weakly admissible formats grows as *N* ^0.78^ to *N* ^1.05^, whereas that of the strongly admissible formats grows as *N* ^0.28^ to *N* ^0.29^. As such, strongly admissible formats are likely better suited for networks of closely packed cells.

## Notes

### Competing Interest Statement

The authors have declared no competing interest.

